# Sensitive inference of alignment-safe intervals from biodiverse protein sequence clusters using EMERALD

**DOI:** 10.1101/2023.01.11.523286

**Authors:** Andreas Grigorjew, Artur Gynter, Fernando H. C. Dias, Benjamin Buchfink, Hajk-Georg Drost, Alexandru I. Tomescu

## Abstract

Sequence alignments are the foundation of life science research, but most innovation focused on optimal alignments, while ignoring information derived from suboptimal solutions. We argue that one optimal alignment per pairwise sequence comparison was a reasonable approximation when dealing with very similar sequences, but is insufficient when exploring the biodiversity of the protein universe at tree-of-life scale. To overcome this limitation, we introduce pairwise alignment-safety to uncover the amino acid positions robustly shared across all suboptimal solutions. We implemented this approach into EMERALD, a dedicated software solution for alignment-safety inference and apply it to 400k sequences from the SwissProt database.

## Introduction

When exploring the diversity of life, we tend to either reduce observations according to similar principles and patterns shared across lineages (comparative method) or aim to deduce the individual mechanistic function with a cause-and-effect-revealing experimental design (functional assessment). In genomics, such attempts translate into comparing genetic sequences according to the similarity of their DNA or protein composition (comparative genomics) or mechanistic analysis of three-dimensional structural conformations of proteins (functional genomics). Recent breakthroughs in protein structure prediction from primary sequence alone [Jumper et al., 2021, Baek et al., 2021] uncovered that integrating comparative and functional genomics into a predictive model can yield groundbreaking insights useful enough to guide mechanistic studies in molecular life sciences. Building on this integrative foundation of sequence comparison and protein structural prediction, we explore how the sequence diversity across the tree of life can be compressed into alignment-robust sequence regions while minimizing the loss of protein structural information.

To compare two biological sequences according to a pre-defined scoring-scheme, pairwise alignment methodologies have proven useful for various practical applications [Needle-man and Wunsch, 1970, Smith and Waterman, 1981]. When constructing a pairwise alignment, a combinatorial space of possible alignment configurations is explored and a single optimal alignment setting is selected (based on the prede-fined scoring-scheme) and reported to equip experimenters with one plausible solution rather than overwhelming them with a wide range of possible solutions. While sufficient for many applications, including protein sequence similarity search, the reduction of comparison to one solution (even when optimal) can cause enormous information loss about biologically relevant, but suboptimal, alignment configurations, thereby systematically biasing the comparative method when applied at tree-of-life scale. One could argue that handling only optimal alignments is the most parsimonious approach to dealing with complexities when scaling to millions or even billions of pairwise sequence comparisons when organizing a diverse sequence space according to their pairwise identities. However, analogous to the concept of point estimates and confidence intervals in statistical parameter inference [Hastie et al., 2009], neglecting the goodness of fit for any application may result in unrealistic technical optima rather than focusing on quantifying the biological relevance (e.g. functional protein configuration) of reported alignment solutions.

It seems therefore surprising that the experimental community has grown accustomed to interpret algorithmically derived optimal alignment solutions as biologically most relevant configuration of pairing similar proteins, although theory clearly states that such approximation may only be reasonable when comparing very similar sequences [Naor and Brutlag, 1994] and not when dealing with distant homologs. We argue that by exploring the space of suboptimal alignment configurations using a quantification method able to capture stable positions across possible alignment paths (*alignment-safe intervals*), novel insights with greater biological relevance can be unveiled and quantified with particular relevance for protein structure evolution at tree-of-life scale.

Fortunately, previously collected evidence suggests that sufficiently screening the suboptimal alignment space for particular configurations that are biologically more relevant can in fact be achieved [Chen and Kihara, 2011, 2008]. Previous work, such as Jaroszewski et al. [2002], Sierk et al. [2010], Cline et al. [2002], has shown that there is a significant connection between the suboptimal alignment space and the structural alignment space and used this link to improve the accuracy of pairwise sequence alignments. It has also been shown that suboptimal alignments are often more accurate than strictly optimal ones and that they contain a high number of correct amino acid residue pairs [Chen and Kihara, 2011], indicating their use in protein structure prediction. Vingron and Argos [1990] showed that so-called *reliable regions*, defined in terms of a *robustness* measure of individual aligned amino acids can identify conserved and functionally relevant regions among two protein sequences. They demonstrate this functional relevance by validating that these conserved regions also correspond to aligned regions of their respective tertiary structures. In detail, Vingron and Argos [1990], Chao et al. [1993] introduce a *robustness* measure for a single pair of aligned symbols between two sequences to assess the difference between the optimal alignment score of the compared sequences without restrictions and the optimal alignment score *not* containing that aligned pair. A related approach suggested by Naor and Brutlag [1994] notes that the space of sub-optimal alignments (whose score is within a difference Δ to the optimal solution) can reveal conserved regions when manually inspecting the “graphic representation” of these possible alignments. While this initial suggestion to explore the suboptimal alignment space yielded promising visual insights, no solution was given on how to automate or scale this approach to millions or billions of pairwise comparisons. Currently, most applications favour multiple sequence alignments (MSAs), Hidden Markov Model based approaches (HMMs), or protein language models (Lin et al. [2022]) when interested in sensitively locating conserved regions inside a set of sequences for large-scale phylogenetic applications (i.e., those positions or regions that remain unchanged in the phylogenetic tree). While providing reliable results in a molecular evolution context, calculating optimal MSAs or constructing relevant HMMs for deep homology searches scales exponentially with the number of sequences, which is computationally expensive and not suitable for tree-of-life applications. In addition, MSAs are designed to process highly similar sequences and often perform poorly when comparing divergent sequences (Penn et al. [2010], Levy Karin et al. [2019]).

Here, we overcome these limitations by introducing the application of *solution safety* [Tomescu and Medvedev, 2017], (Methods - Definition 1), for pairwise protein sequence alignments. With alignment safety we can explore the space of optimal *and* suboptimal alignment configurations (i.e. possible pairings of amino acids), and find entire intervals that are common to all or to a given proportion of alignment solutions. We implement this approach in a command line tool, EMERALD [Grigorjew et al., 2023]. Instead of forcing two (possibly very diverse) sequences into a single optimal alignment configuration, EMERALD embraces the diversity of possible alignment solutions, by revealing *alignment-safe intervals* of the two sequences which appear as conserved (and not even necessarily identical) in the entire space of optimal and suboptimal alignments (Figure 1). To demonstrate the effectiveness of this procedure for sizeable protein comparisons, we first cluster all protein sequences stored in the Swiss-Prot database [Consortium, 2021, Bairoch and Apweiler, 1997] using DIAMOND DeepClust [Buchfink et al., 2021] (Results and Methods) and apply EMERALD to each non-singleton cluster. With this comprehensive analysis, we aim to explore whether our method can provide a competitive solution to project alignment-safe primary-sequence intervals onto the structural conformation of proteins and thereby accelerate the biologically relevant exploration of sequence evolution and their corresponding structural divergence across the tree of life.

**Figure 1:**
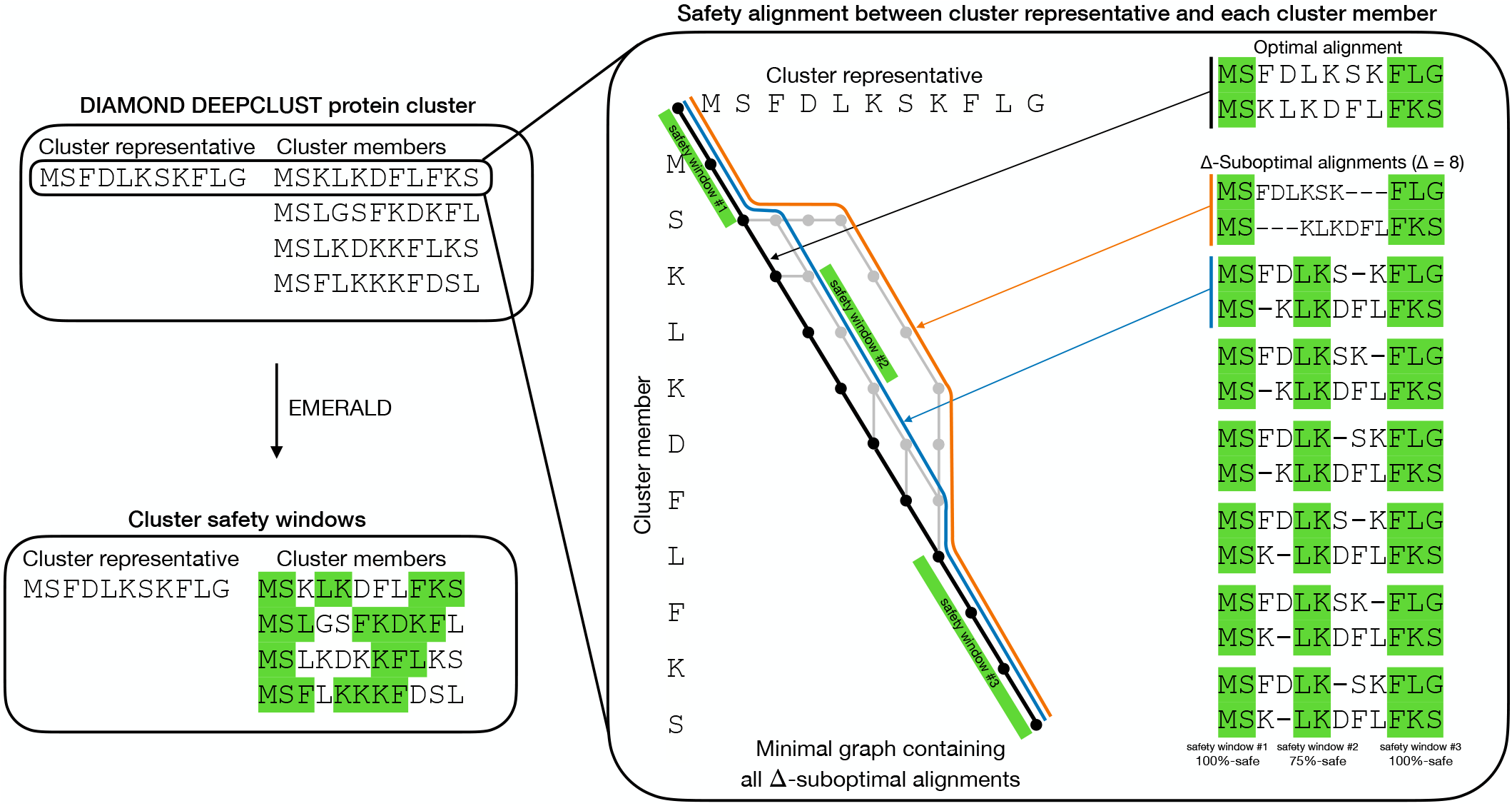
Schematic representation of EMERALD’s safety window calculation of a DIAMOND DeepClust cluster containing 4 member sequences. EMERALD performs a pairwise global alignment between the cluster representative against each of the 4 cluster member sequences using affine gap costs and BLOSUM62 as substitution matrix. For the first sequence pair, the right-hand side illustrates the suboptimal alignment graph and their corresponding suboptimal alignment configurations between the two sequences listed as Δ*-suboptimal* alignments (An alignment is Δ*-suboptimal* if its score is not more than Δ smaller than the optimal score). The illustrated graph is one of minimum size to fulfil the property of including all Δ-suboptimal alignments [Naor and Brutlag, 1994] (here, we choose Δ = 8). Source-to-sink paths in the graph correspond to suboptimal alignments; nodes and edges on the unique optimal alignment path are shown in black, while those configurations on a Δ-suboptimal path are illustrated in gray. The optimal alignment path is colour coded in black and the two top Δ*-suboptimal* alignment paths illustrated in orange and blue. For *α* = 0.75 and Δ = 8 we obtain three safety windows shown as green intervals. These three coloured safety windows correspond to subpaths contained in at least *α* = 0.75 (i.e. 75%) of all source-to-sink paths (i.e., of all Δ-suboptimal alignments). Note that the middle safety window is not captured (i.e., contained) by the (unique) optimal alignment, in black, and is only revealed by the subgraph of all Δ-suboptimal alignments. Finally, we project the safety windows onto the cluster member (and cluster representative sequence) as explained in (Methods). This procedure is repeated for all possible pairwise comparisons between the representative sequence and the 4 members, thereby obtaining (*α*, Δ)-safety windows for each cluster member (bottom left).

## Results

The main purpose of this study is to notify the genomics community about the advantages of exploring the suboptimal alignment space when dealing with comparisons of vastly divergent sequences where subsequent structural information will be used to make claims about protein sequence and fold evolution at tree-of-life scale. EMERALD allows to achieve this task by equipping users with an automated and scalable software solution to infer alignment-safe subsequences extracted from the suboptimal alignment neighborhood (Figure 1), which we demonstrate experimentally to correspond to conserved regions of the underlying protein structure (such as alpha-helix, etc).

### Defining Alignment-Safety for Pairwise Protein Sequence Alignments

Initially developed in the context of genome assembly [Tomescu and Medvedev, 2017], and later extended to other applications admitting multiple solutions, such as flow decompositions for RNA transcript assembly [Khan et al., 2022], and RNA folding [Kiirala et al., 2019], *solution safety* identifies a set of partial solutions (e.g., an interval of an alignment) to infer *safe* positions or windows present in all optimal and Δ-suboptimal configurations to the problem. Using this definition, previous work by Vingron and Argos [1990], Chao et al. [1993], Naor and Brutlag [1994] can be reintroduced in the context of studying maximally safe partial alignments, whereby *maximal* denotes the property that such partial alignment cannot be extended left or right without losing the characteristic of being safe.

To achieve this property at scale and with biological relevance, we introduce novel computational aspects of the alignment-safety concept and implement these into the *C++* command line tool EMERALD (Methods). First, we generalize the notion of safety by defining a partial solution to be *α-safe* (*α*∈ (0, 1]) if it is present in at least a proportion *α* of all possible solutions. We further generalize *α*-safety to the space of Δ-suboptimal solutions from [Naor and Brutlag, 1994] that are at most Δ away from the optimal solution, by defining a joint-parameter (*α*, Δ) which captures partial solutions that appear in at least a proportion *α* of all Δ-suboptimal solutions. In other words, while Δ captures the boundaries of the suboptimal alignment space that a user wishes to explore, *α* regulates the quantile-range over all possible alignment solutions. The joint-parameter (*α*, Δ) then allows users to specify the suboptimal alignment space within a particular quantile range over all solutions that shall be explored. This approach allows us to address the fact that an (*α*, Δ)-safe partial solution is an interval of arbitrary alignment length (not only a single pair of aligned symbols, as is the case for the robustness measure from [Vingron and Argos, 1990]). Together, we denote all such (*α*, Δ)-safe intervals as the collection of *alignment-safe protein sequence intervals* or in short *safety windows*. Intuitively, Δ allows sufficient exploration across the suboptimal alignment space within an *α* range, thus enlarging the solution space, and leading to shorter safety-windows, while *α* relaxes the safety requirement by enforcing that only a *α*-fraction of the Δ-suboptimal alignment-configurations need to have the same amino acid to extend safety windows. We show that optimizing the *α* and Δ configuration can be a powerful tool for regulating the biological relevance safety-windows can capture in diverse protein sequences and their respective three-dimensional structures.

Additionally, we explore the biological signatures that can be captured when exploring the suboptimal alignment space using our alignment-safety approach. In order to annotate which biological features alignment-safe residues may encode in a particular threshold- and substitution-matrix configuration, we use the Stride [Frishman and Argos, 1995] annotation such as alpha-helices and loops to exemplify how users can biologically assess the output of EMERALD. However, we would like to point out that users need to pay close attention to what suffices as valid ‘ground truth’ for their bench-marking, since EMERALD is agnostic to particular biological applications and the choice of substitution matrix may induce a well characterized amino acid bias for evolutionary more stable residues [Notredame et al., 2000, Chatzou et al., 2016, Baltzis et al., 2022]. Recent advancements in protein structure prediction can also provide users with a more structurally oriented validation dataset [Baltzis et al., 2022]. While it is well understood that secondary structure elements align better than non-secondary structure elements or disordered parts of proteins, our methodology can reveal the concrete regions of a protein sequence (alignment-safety windows) that are the same across all (considered) suboptimal alignment solutions. This feature of EMERALD allows users to explore how robust certain sequence regions are to alternative alignment configurations and different substitution matrices. For this purpose, we extracted the secondary structure from the Swiss-Prot database for each protein sequence using the command line tool Stride [Frishman and Argos, 1995]. For each residue in a sequence, Stride assigns a secondary structure type, in our case: “AlphaHelix”, “310Helix”, “Pi-Helix”, “Strand”, “Bridge”, “Coil” and “Turn”. We further distinguish secondary structure types by placing them into two distinct categories: stable and not stable. Coil, Strand and Turn were labeled as not stable while the rest of the secondary structure types were labeled as stable. While this classification is naive in the first instance (for example, because there are well-known cases where Coils, Strands and Turns can be stable to fix certain protein conformations and vice-versa, alpha-helices can be fairly flexible and unstable (Levy Karin et al. [2019], Bondos et al. [2021])), we use this distinction only to exemplify how users can categorize known protein-structural features into distinctive structural feature classes to benchmark the suboptimal alignment-space for their domain-specific application and quantification of biological relevance. The main motivation behind such categorization is to test which regions of a protein structure are robustly encoded in alignment-safe intervals. We envision that users make extensive use of this benchmarking setup to explore, quantify and test the biological relevance of harnessing the suboptimal alignment-space for encoding structural features and protein evolution of their (divergent) sequences of interest.

### *In silico* experimental design to investigate the biological relevance of alternative suboptimal alignment configurations

The concept of solution-safety [Tomescu and Medvedev, 2017] guarantees the following property. If the true alignment-configuration is inside the full solution space of the dynamic programming matrix (i.e., optimal and Δ-suboptimal alignment-configurations), then windows that are the same across all such alignment-configurations (safety windows) are also part of the true alignment (which cannot be observed in reality). In other words, analogous to sampling theory in statistics where the ground truth of a data universe cannot be measured, we assume that a representative sample drawn from this data universe allows us to infer rules and principles about the data universe itself. Alignment-safety windows can thus be seen as analogy of a representative sample drawn from the true (unobserved) alignment. The biological relevance of this approach can then be studied by annotating these alignment-safe windows/positions in regard to their overlap with protein secondary structural features, their contribution to protein selection pressures such as synonymous vs nonsynonymous substitution rates (dN/dS), and mappings onto AlphaFold2 structures or their underlying multiple-sequence alignments (MSAs). The focus of our experimental design is therefore, to test whether exploring the suboptimal alignment space when dealing with millions or billions of pairwise alignments spanning a comprehensive sequence diversity across the tree of life yields sufficiently more information (compared to a single optimal alignment configuration) to increase biologically relevant inference power when dealing with protein structure prediction and structural evolution tasks. To achieve this, we retrieved the Swiss-Prot database from [Bairoch and Apweiler, 1997] (June 2021), which represents a manually curated subset of the UniProtKB [Consortium, 2021] database. In the version of June 2021, Swiss-Prot contains approximately 560k protein sequences which can be retrieved as a FASTA-file. After data retrieval, we filter out protein sequences that did not have corresponding AlphaFold2 predicted three-dimensional structures [Jumper et al., 2021, Varadi et al., 2022]. We then clustered this dataset with DIAMOND DeepClust, a new sensitive deep-sequence-clustering method implemented into DIAMOND since version 2.1.0 [Buchfink et al., 2021]. We filter out cluster members with no stable bases and clusters of size 1, resulting in 15934 clusters and 396k sequences in total. To achieve a comprehensive overview of deep-homology associations between proteins across the full diversity intrinsic to the 396k Swiss-Prot sequences, clustering was carried out using a percent-identity threshold of 20% and 75% length coverage. This threshold was motivated mainly by two factors: 1) conformity with the twilight zone of protein evolution where sequence identity greater than 20-35% can still be reliably associated with structural similarity above this zone, while this association is ‘breaking’ otherwise [Rost, 1999]; and 2) to maximize the diversity of the protein sequence space across the tree of life. It is important to note that although the minimum pairwise identity threshold ensures that distant similarities are detected, some clusters can yield pairwise sequence compositions that have significantly higher identity-relations than the chosen threshold of 20%, since DIAMOND uses no upper identity threshold when clustering.

Next, we run EMERALD to calculate safety windows of all sequence and representative pairs of each cluster. To explore the influence of the suboptimal alignment space depth on capturing biologically relevant features, we test various parameter-combinations performed using three different *α* parameters (0.51, 0.75, 1) and seven Δ parameters (0, 2, 4, 6, 8, 10, 15), values that are in the same order of magnitude as the BLOSUM62 metric [Henikoff and Henikoff, 1992], resulting in 21 safety window calculations per cluster (Figures 2 to 3). As cluster representative, we selected the cluster centroids reported by DIAMOND DeepClust’s greedy set cover method. Finally, we benchmarked the CPU runtime and maximal memory consumption of these runs (Figure 4).

**Figure 2:**
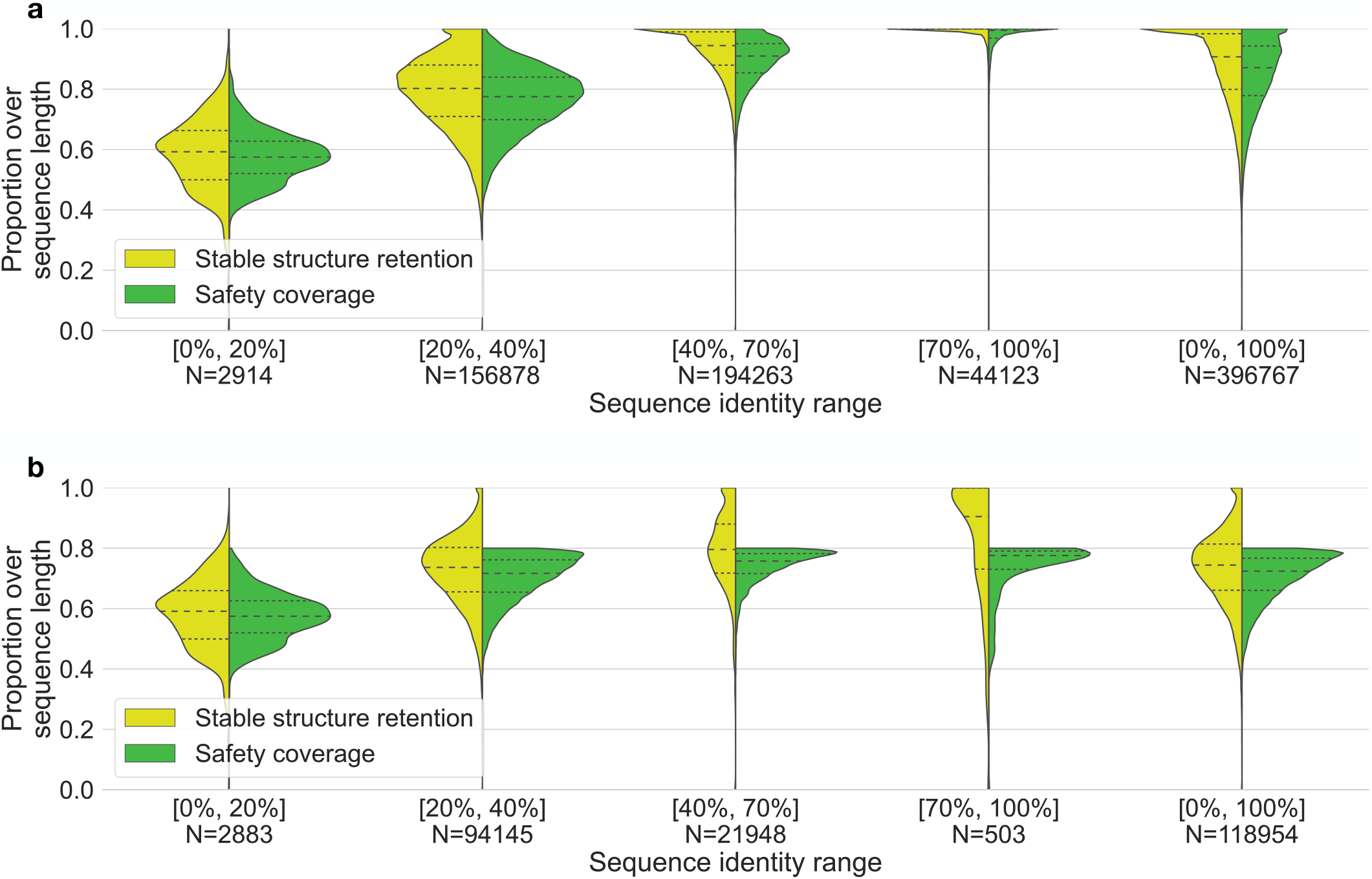
Comparing stable structure retention and safety coverage (y-axis) for several sequence identity ranges between cluster members against the cluster representative (x-axis) carried out with EMERALD parameters *α* = 0.75 and Δ = 8 on all obtained sequences, where *N* denotes the number of sequences in each identity range. The dashed lines indicate medians and the dotted lines the first and third quartiles. The safety coverage increases with higher identity ranges due to the smaller size of the suboptimal alignment space for high identity sequence pairs. Since the stable structure retention quantifies the proportion of stable amino acids that are alignment-safe, it increases alongside an increase in safety coverage. The high stable structure retention results indicate that EMERALD is indeed able to capture biologically relevant stable positions, as they make up a higher proportion when restricting the sequence to safety windows. **a** All sequences are included (i.e., with safety coverage of at most 100%). **b** Sequences restricted to those having safety coverage of maximum 80%.

**Figure 3:**
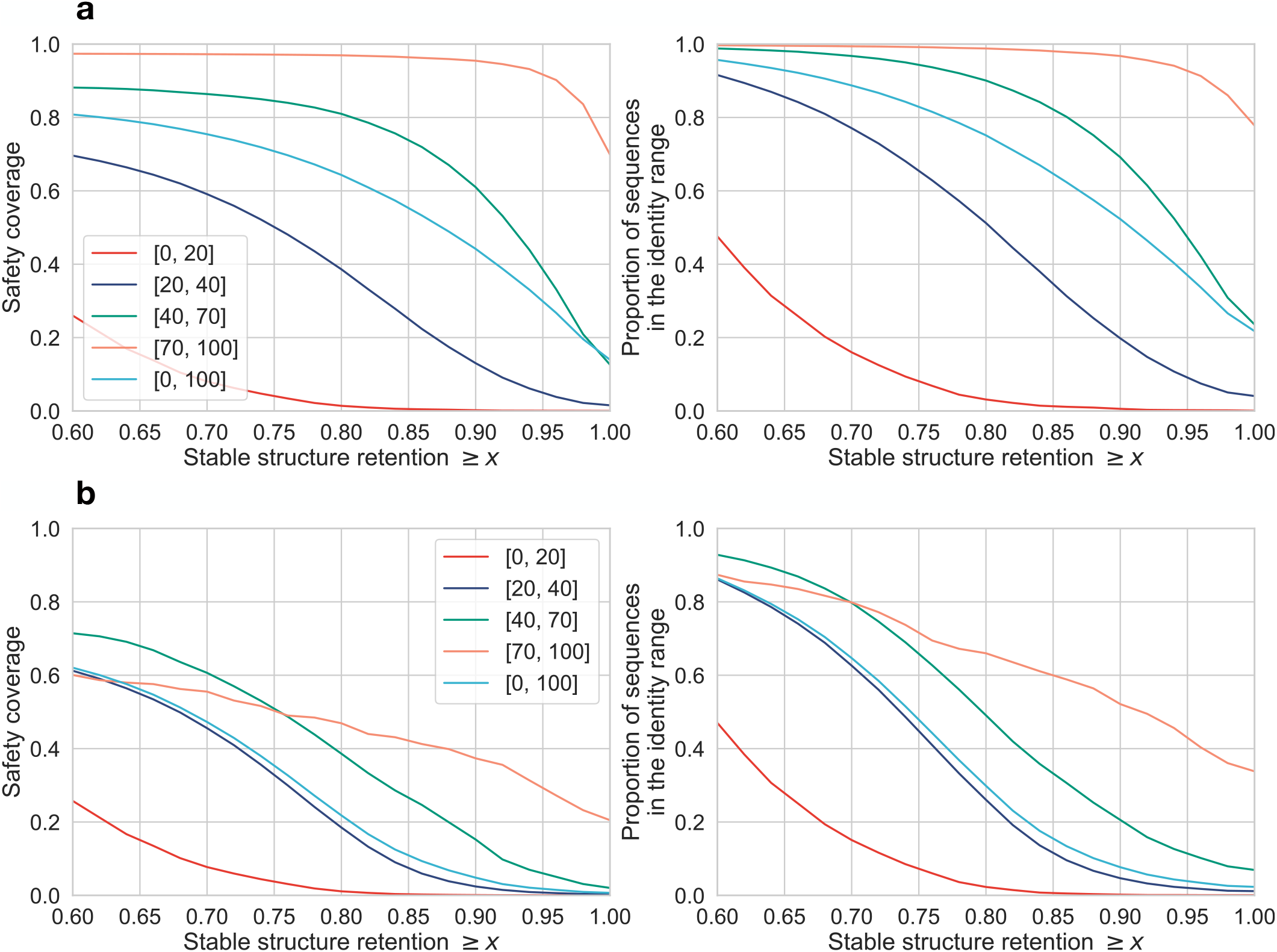
For a given threshold *x*, safety coverage of the sequences whose stable structure retention is at least *x* (on the left), and the proportion of the sequences whose stable structure retention is at least *x* (on the right). The results are split in the identity ranges from Figure 2. **a** All 396k sequences are included. **b** Restricted to sequences whose safety coverage is at most 80%. The turquoise curves, which cover sequences of all identities, show in **a** that around 20% of the sequences (right plot) have a stable structure retention of 100% and only 15% of safety coverage (left plot), while the green curves, which cover sequences from the identity range 40% - 70%, show in **b** that 10% out of the sequences with at most 80% safety coverage inside this identity range (right plot) have a stable structure retention of 100% and only a safety coverage of around 5% (left plot).

**Figure 4:**
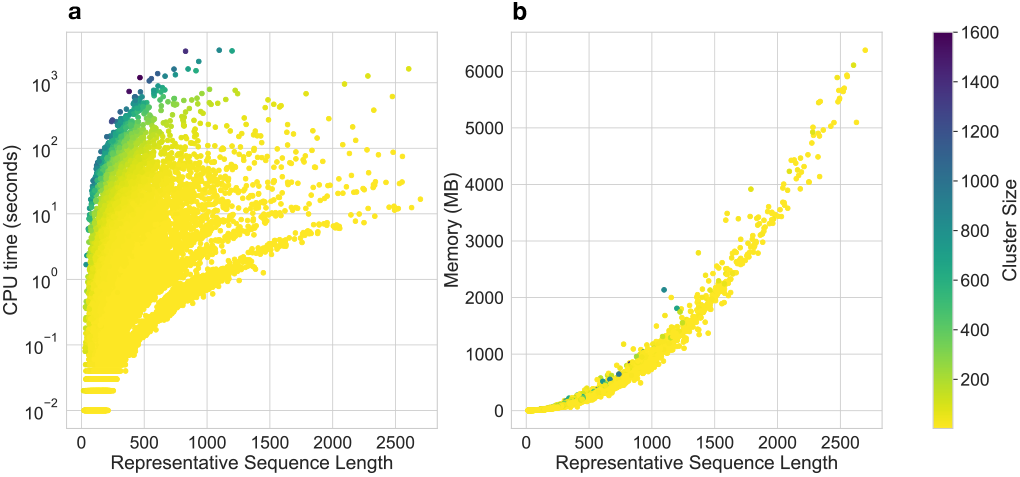
CPU runtime of EMERALD. (**a**) Computational run time for all DIAMOND DeepClust clusters generated from the filtered Swiss-Prot database using the threshold combination *α* = 0.75 and Δ = 8 and calculated on a single thread. Each dot corresponds to a protein sequence cluster and the colour of each dot indicates its number of corresponding member sequences. (**b**) Maximum memory consumption of EMERALD runs. For all trialed *α* and Δ parameter settings the average memory consumption for each cluster ranged between 205 − 207 Mb and the average run time between 11.4 − 11.5 seconds.

**Figure 5:**
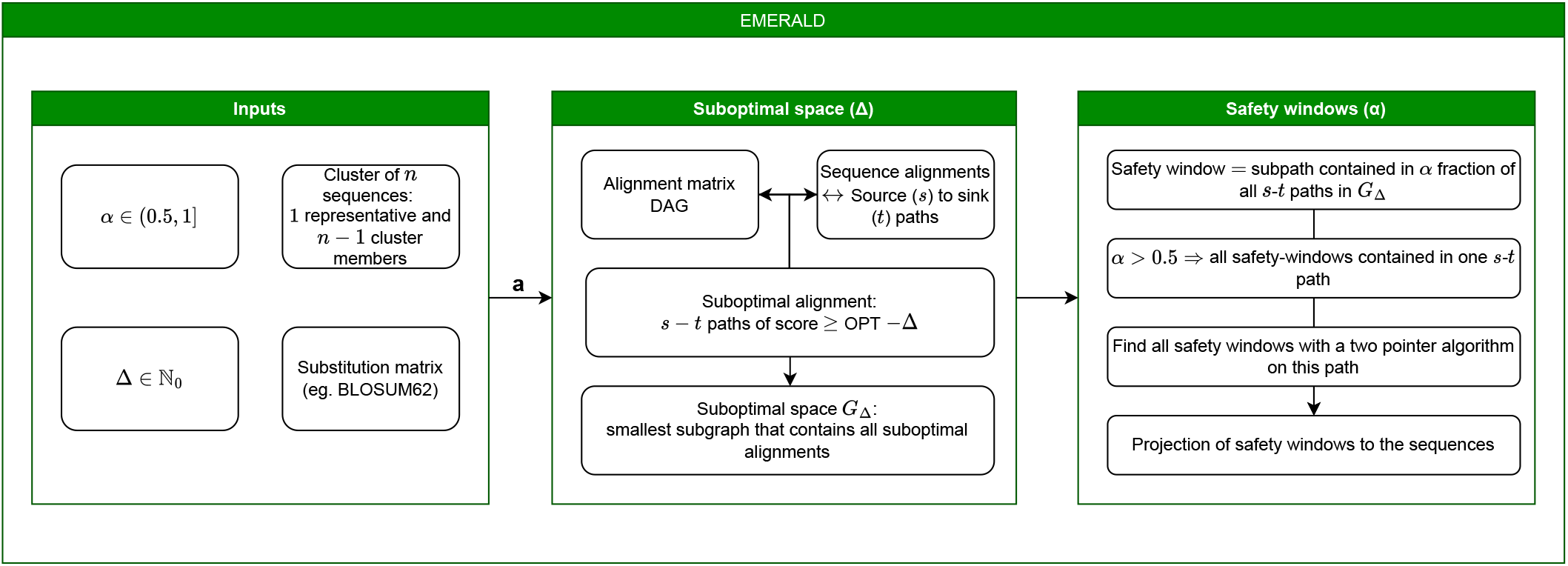
Conceptual overview of EMERALD’s safety window calculation workflow. As input EMERALD receives a set of clusters in *fasta* format. For example, such protein sequence clusters can be generated using DIAMOND DeepClust or alternative clustering methods. Next, users can specify the scoring matrix (e.g. BLOSUM62) according to which optimal alignment configurations will be determined. Each cluster member sequence is then globally aligned against the cluster representative sequence (centroid) using the pairwise Needleman-Wunsch alignment algorithm. The resulting dynamic programming (DP) matrix of each pairwise comparison is then encoded as a graph data structure to search for optimal and suboptimal alignment paths according to the selected scoring matrix and the threshold configurations defining the suboptimal alignment space. Once all alignment-safe intervals are computed, EMERALD projects these safety intervals (safety windows) back to the representative sequence, thereby annotating the sequence intervals that are robust across all possible alignment configurations within the suboptimal alignment space.

### Benchmarking and analysis of stable versus unstable protein structural features that are encoded by alignment-safe windows

For quantification and comparison of safe intervals in the context of stable versus unstable bases, we utilise several benchmarking metrics: safety coverage, stable coverage, stable structure overlap and stable structure retention. Safety coverage is defined by the portion (%) of a protein sequence which is reported by EMERALD as (alignment-) safe, while stable coverage is the portion (%) of a sequence that is considered stable according to the distinct protein-structural features classification (user-defined) described in the previous section. We further classify a sequence position as *true positive (TP)* if it is both safe and stable, *false negative (FN)*, if it is not safe but stable, and *false positive (FP)* if it is safe but not stable. Based on these distinctions, we compute the metrics *stable structure retention (recall) = TP/(TP+FN)*, indicating the percentage of stable positions captured by safety windows, *stable structure overlap (precision) = TP/(TP+FP)*, indicating the proportion of safety windows that is also stable, and their combination, *F1-score*, which is the harmonic mean of the stable structure retention and overlap. This analytics tool allows users to benchmark according to their own distinction of structural classes when determining how much information is gained when incorporating the suboptimal alignment space to the sequence comparison task.

We represent the sequence identity ranges as intervals of real numbers with notation [*i, j*] where *i* defines the lower identity boundary and *j* the upper identity boundary. Figure 2 illustrates the stable structure retention and safety coverage benchmark for our 396k sequence dataset. Naively, we expect the stable structure retention and the safety coverage to be similar, as we are likely to cover e.g. 60% of all stable elements if 60% of the sequence is covered. Similarly, if all amino acids are alignment-safe, the stable structure retention is equal to 100% meaning that no further biological information can be obtained. Thus, we are usually interested in an increased gap between the stable structure retention and the safety coverage. Ideally, the stable structure retention would always be close to 100% with all varying safety coverages, meaning that safety corresponds perfectly to stable elements. Figure 2 further shows that as we consider sequences that are more dissimilar, safety coverage decreases, since we increase the space of optimal and suboptimal solutions. Figure 2 **b** shows the stable structure retention that is restricted to all sequences with a safety coverage of at most 70%, resulting in a decrease of the stable structure retention. In both, Figure 2 **a** and **b**, the average values of the stable structure retention exceeds the average value of the corresponding safety coverage, and in the identity range [40%, 70%] the average stable structure retention is at 80% even when we restrict the sequences to have no more than 80% safety coverage. This result shows that the proportion of stable positions inside the safety windows is larger than the proportion of stable positions out of all the sequence (i.e., (stable *∩*safe)*/*safe *>* stable*/*sequence length). In addition to Figure 2, we also assessed the safety coverage of each individual amino acid structure type in Supplementary Figure 2, which shows that unstable amino acids have a smaller safety coverage than stable ones. “Coil” has the smallest coverage followed by “Turn”, with an exception of “Strand” having a higher coverage than “AlphaHelix”, which we defined as stable. EMERALD sufficiently captures amino acids of type “Bridge” and “310Helix”, despite STRIDE only assigning these types to 0.8% and 3% of all amino acids, respectively.

Figure 2 **a** illustrates that the best tradeoff between safety coverage and stable structure retention is the identity range [40%, 70%] sequence identity. This range setting introduces the biggest gap between medians while ensuring that the stable structure retention does not drop below 70%. In Figure 2 **b** the median of the stable structure retention in the identity range of 40%-70% is at 80%, despite the safety coverage being restricted to be at most 80%. Furthermore, we analysed the stable structure retention over all combinations of parameters (Supplementary Figures 3 to 5). Overall, the stable structure retention is always slightly larger than the safety coverage. For Δ = 0 (i.e. when we consider only optimal alignment parts) and all three *α* values 0.51, 0.75 and 1.0, the safety coverage is close to 100% for identities of at least 20%, which implies that the substitution matrix is very good at finding a unique optimal alignment, which motivates exploring the suboptimal alignment space. As we predicted, decreasing *α* corresponds to increasing the stable structure retention and safety coverage, while increasing Δ corresponds to decreasing them.

The choice of parameters also has an impact on the contiguity of safety window lengths and counts. For example, when selecting very conservative parameter settings, a larger proportion of shorter safety windows are generated. To buffer this effect, we added the additional constraint as a new EMERALD parameter –m (–windowmerge) which determines whether short safety windows that appear in close proximity (i.e., intersecting or adjacent to each other), are subsequently joined into one large safety window. We quantify this parameter selection behaviour in Supplementary Table 1, which now provides the average sequence and safety window lengths (summary statistic of contiguous stable intervals). As expected, lowering the pairwise sequence identity results in shorter and overall less contiguous safety windows. Additionally, a substantial amount of the safety windows of sequence alignments in the [70%, 100%] identity range have a length exceeding 50% of their full sequence length. Over the whole dataset and all combinations of parameters, the stable structure overlap stays constant (Supplementary Figure 1) with the median at around 43%.

Figure 3 further explores the relationship between stable structure retention and safety coverage. It restricts the sequences in the *x*-axis to those with stable structure retention of at least *x*, plotting the safety coverage (on the left) and the proportion of considered sequences (on the right). Strikingly, in Figure 3 **a** we can see that for about 20% of all sequences, or of all sequences in the identity range [40%, 70%] (turquoise, and green curves, respectively on the right for *x* = 1.00) EMERALD retains *all* their stable positions, with a safety coverage of only 15%. This illustrates that, in contrast to optimal alignment approaches, for lower identity bounds (dissimilar sequences) EMERALD manages to reveal structurally conserved intervals. Similarly, EMERALD achieves a stable structure retention of 80% for around 75% of all the sequences with a safety coverage of around 65%. In other words, EMERALD declares 65% of the sequence as safe, capturing 80% of all stable amino acids. This result further shows that by relaxing the stable structure retention criteria from 100% down to 80%, EMERALD is able to reduce the sequences to their safety intervals from full length to 65%. In Figure 3 **b**, we analogously restrict the sequences only to those of safety coverage of at most 80%. Here, for example, the green curve shows that around 10% of the sequences in the identity range 40% - 70% can be reduced to a safety coverage of nearly 5%, without losing the stable structural retention constraint of 100%. This suggests that EMERALD embraces the sequence diversity to narrow down the structurally-conserved context of a sequence.

Supplementary Figure 6 analyses the safety coverage and the F1 score, and it shows that, as we consider clusters with smaller post-computed identity values, F1-score has only a minor decrease, but safety coverage has a marked decrease. This indicates additionally that safety windows have a better ability to capture stable structural elements in clusters with smaller identity values.

Finally, we benchmarked EMERALD’s run time and memory consumption (Figure 4) on a computing server with 32 cores (2x hyper-threading, 64 threads) and 512GB of RAM using the clustered 396k sequences from the Swiss-Prot database. Overall, none of our clusters ran more than 17 minutes, and, except for two clusters, they did not use more than 6.5GB of memory. While the time increases quadratically in the lengths of sequences, trivially it increases only linearly with the number of sequences in the cluster. This result illustrates that EMERALD can scale to millions of sequences and thousands of clusters to explore the protein universe across the tree of life.

## Discussion

For the past decades pairwise sequence alignments have served silently and reliably as foundation of comparative and functional genomics applications. Algorithmic innovation focused on computing ever faster heuristics for retrieving optimal alignment solutions at scale or extending comparisons to multiple sequence alignments or Hidden Markov Model based statistical alignment approaches. However, only little innovation occurred in the pairwise alignment field on quantifying the biologically most relevant alignment configuration from a collection of up to millions of possible (suboptimal) alignment solutions. While biological meaning in the context of alignment optimization is a vague concept, in the early days of comparative and functional genomics the ability to encode structural information of proteins was among the main applications underlying the benchmarking of biologically meaningful alignment configurations [Chen and Kihara, 2011, Hastie et al., 2009]. The consensus then was that focusing on optimal alignments is a reasonable heuristic when dealing with highly similar sequences, since the optimal alignment solution can indeed encode a good representation of conserved protein structure such as alphahelix (Ranwez and Chantret [2020]). Still applied as a main assumption in functional genomics today when employing pairwise alignment searches, protein sequences are usually screened for high similarity across distant species or strains to test whether this retained sequence identity translates into structural conservation and potential functional similarity.

In this study, we aimed to determine whether quantifying suboptimal alignment configurations in pairwise sequence alignments can significantly improve the sensitivity of identifying relationships between features of protein structure across the tree of life when analyzing a large diversity of protein sequence space and making comparisons between hundreds of thousands of species [Buchfink et al., 2021]. To approach this quest, we designed an *in silico* experiment to annotate all alignment-safe positions of the Swiss-Prot database. Using this annotation approach, we investigate how the quantification of the suboptimal alignment space can refine biologically relevant interpretations such as conserved protein structural features. We asked how information about suboptimal alignment configurations at scale can be harnessed to predict protein structural change when the proportion of alignment-safe positions in distant sequence alignments is reduced. Finally, we introduced *alignment-safety* as a new methodology to approach such questions and the command line tool EMERALD to implement our methodology at scale.

While previous work focused on assessing the quality or statistical robustness of optimal alignment configurations in comparison with a set of suboptimal alignment solutions (optimal alignment neighborhood) [Kschischo and Lässig, 1999, Schlosshauer and Ohlsson, 2002, Zhang and Marr, 1995] in the context of protein homology modeling or threading, our work significantly extends these concepts to associate protein sequence evolution with their respective structural change while scaling to millions of pairwise comparisons and trillions of suboptimal solutions at tree-of-life scale. We achieve this by inferring the robust amino acid positions across a range of suboptimal alignment-configurations. As a result, we learned that when attempting to relate the sequence biodiversity of the protein universe to the evolution of protein structure, a detailed inference and exploration of how alignment-safe positions are retained or lost throughout the tree of life can serve as robust proxy for how conserved structural features change over evolutionary time. We interpret this result such that alignment-safe positions can be associated with the stably folded backbone of a protein structure and that further research is required to unveil the causal associations between alignment-safe positions and positions giving predictive signatures when inferring structural conservation above the twilight zone of protein evolution [Rost, 1999].

### Conclusions

We envision that studies exploring the suboptimal alignment space when comparing protein sequences pairwise across the protein universe will stimulate further research attempting to address the remaining shortcomings of fold predictions such as dealing with disordered parts of proteins and incorporating diverse mutation patterns into fold evolution predictions. Our alignment-safety methodology and EMERALD software are designed to assist these efforts of associating new information gained from suboptimal alignments with biologically relevant phenotypes of proteins in an evolutionary context. We designed our approach to be sufficiently fast and sensitive to scale to millions of sequences with various degrees of divergence to supply the data-intensive demands of the biosphere genomics era.

## Methods

Conceptually, we build on the approach introduced by Naor and Brutlag [1994] and vastly extend its methodological depth and computational scalability for tree-of-life scale applications. In detail, the approach introduced by Naor and Brutlag [1994] is restricted to consider only aligned symbols that are part of *all* suboptimal alignments, ignoring pairs which are part of the optimal alignment itself. For example, suppose that an aligned pair is common to all the 1000 optimal alignments of score *q*, there is a single suboptimal alignment of score *q*− 1 not containing the pair, and the next suboptimal alignments not containing the pair have score *q* −100. In this case, the single suboptimal alignment of score *q* −1 makes the pair “not conserved” under the approach of Naor and Brutlag [1994], and drastically decreases the robustness of the pair (from 100 to just 1), since the robustness measure cannot quantify the *proportion* of (sub-)alignments containing the pair. In addition, they define robustness independently for each aligned amino-acid pair thereby excluding the opportunity to quantify robust regions, which are often observed in natural settings. As a result, although Naor and Brutlag [1994] provide a first theoretical template to study suboptimal alignment spaces, they fail to deliver scalable algorithmic solutions, software tools, and biological validation to investigate the protein sequence diversity space in the context of robust - alignment safe regions for tree-of-life scale applications. In fact, their methodology does not exceed a manual analysis of graphical alignment representations with little potential for automation and efficient scaling.

### Inference of Alignment-Safe Protein Sequence Windows

We formally introduce the calculation of alignment-safe intervals through the thresholded exploration of the suboptimal pairwise alignment space. Let *A* and *B* be two strings over an alphabet ∑ of length *n* and *m*, respectively. In this study, we refer to an optimal alignment between *A* and *B*, when a particular alignment *maximizes* a scoring function based on an externally provided match/mismatch cost matrix. First, we consider scores provided by a amino acid substitution matrix (such as BLOSUM62 [Henikoff and Henikoff, 1992]) and solely focus on optimal alignments, before we introduce affine-linear gap costs in the subsection Introducing gap penalties [Myers and Miller, 1988] and the suboptimal alignment space to explain the necessary changes in our approach.

In sequence bioinformatics it is established that such optimal global alignments between *A* and *B* can be computed via dynamic programming [Needleman and Wunsch, 1970]. A common result is that alignments of maximum score are in bijection with maximum-weight (*optimal*) paths in the directed acyclic graph (*DAG*) which corresponds to this dynamic programming table. More formally, we can define the *alignment DAG of A and B* as *G*(*A, B*) = (*V, E*), with *V* = *{*0, …, *n}× {*0, …, *m}*, with each node (*x, y*) denoting three out-going edges in *E* to (*x* + 1, *y*), (*x, y* + 1) and (*x* + 1, *y* + 1) (and thus, the DAG has unique source *s* = (0, 0) and unique sink *t* = (*n, m*)). We denote paths in a graph by the notation *P* = (*v*_1_, *v*_2_, …, *v*_*k*_) ∈ *V* ^*k*^ and their restriction by *P* [*v*_*L*_..*v*_*R*_] = (*v*_*L*_, *v*_*L*+1_, …, *v*_*R*_). The first two of the edges correspond to a gap and the third edge corresponds to an alignment of the symbols *A*[*x*] and *B*[*y*]. If we assign the scores of the substitution matrix as well as the gap score to the corresponding edges as weights, then finding an optimal alignment corresponds to finding a maximum-weight path from *s* to *t* (*s-t path*) in *G*. We define such score maximizing paths as *optimal*. We are then interested in discovering those *safe* partial alignments that are common to all optimal alignments.

We can further generalize the notion of safety by also considering a parameter *α*∈ [0, 1] and analogously explore *α*-safe partial alignments that appear in at least the proportion *α* of all optimal alignments. For *α* = 1, 1-safety coincides with safety, and considering *α <* 1 allows us to have potentially longer *α*-safe paths. We can define these notions formally based on the graph-centric definition of an optimal global alignment.

#### Definition 1

Let *α*∈ [0, 1], let *A* and *B* be two strings, and let *G*(*A, B*) be the global alignment DAG connecting *A* and *B*. We denote that a path *P* in *G* is

- *safe*, if *P* is a subpath of all optimal *s*-*t* paths of *G*;
- *α-safe*, if *P* is a subpath of at least an *α* proportion of all optimal *s*-*t* paths of *G*.
- *maximally α-safe*, if it is not a subpath of a longer *α*- safe path.

We define the set of all maximal *α*-safe paths as *α-safety windows*. We will omit *α* when it is clear from the context.

Given the alignment DAG *G*(*A, B*) that connects *A* and *B*, we define *G*_0_(*A, B*) = (*V*_0_, *E*_0_) as the unweighted minimal subgraph of *G*(*A, B*) to include all optimal *s*-*t* paths in *G*(*A, B*) (thus *s* and *t* are nodes in both, *G*(*A, B*) and *G*_0_(*A, B*)). Backtracing an optimal solution in *G*(*A, B*) can be done by taking any in-coming edges of any node (*x, y*) that is part of an optimal path. Due to the bijection property of *G*_0_, we can ignore edge weights and focus on exploring only the set of all its *s*-*t* paths, since these are bijectively linked with the optimal alignments between *A* and *B*. We note that *G*_0_(*A, B*) is weakly connected (i.e., there is an undirected path between any two pairs of nodes, since by definition every node appears in some *s*-*t* path of *G*_0_(*A, B*)). It is easy to notice that the safe edges of *G*(*A, B*) are exactly the edges that when removed from *G*_0_(*A, B*) leaves *G*_0_(*A, B*) no longer sparsely connected. Such edges are also referred to as *bridges* of the undirected graph underlying *G*_0_(*A, B*) and they can be computed in linear time and in proportion to the size of *G*_0_(*A, B*) [Tarjan, 1974].

Here, we propose a refined approach to compute *α*-safe paths based on counting *s*-*t* paths in *G*_0_(*A, B*). For each node *v*, let *d*(*v*) be the number of paths from *v* to *t*. We can calculate these numbers with the following recurrence:

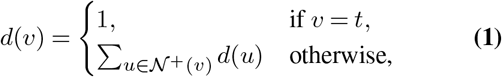

where 𝒩^+^(*v*) = *{u*∈ *V*_0_ | (*v, u*) ∈ *E*_0_ *}*. Since *G*_0_ is a DAG, the recurrence is well-defined, and it stores the total number of *s*-*t* paths in *d*(*s*). Analogously, we can also compute the number *d*_*r*_(*u*) of *s*-*u* paths for any *u* ∈ *V*_0_. Given these two counts for any edge *e* = (*u, v*) ∈ *E*_0_, we can define

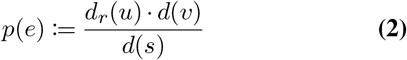

as the proportion of all *s*-*t* paths of *G*_0_ that *e* is part of. We can also understand *p*(*e*) as the probability of *e* appearing in an arbitrary *s*-*t* path of *G*_0_. Likewise, given a path *P* = (*v*_1_, …, *v*_*k*_) of nodes *v*_*i*_ ∈ *V*_0_, the proportion of *s*-*t* paths that *P* is part of is given by

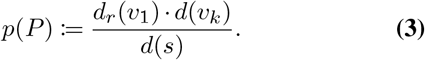

Thus, *P* is *α*-safe if and only if *p*(*P*) ≥ *α*.

The following lemma shows that if *α >* 0.5, then there exists an *s*-*t* path *P* ^*\**^, such that any *α*-safe path is a subpath of *P* ^*\**^. This not only simplifies the algorithm, but it also makes it computationally efficient since it guarantees that the number of maximal *α*-safe paths is proportional to the size of this path (i.e., 𝒪 (*n* + *m*)). Moreover, since *P* ^*\**^ is an *s*-*t* path in *G*_0_, it corresponds to an optimal alignment between *A* and *B*, and thus all safety windows can be reported as intervals of this alignment.

#### Lemma 1

Let *A* and *B* be two strings based on an amino acid alphabet ∑. If *α*∈ (0.5, 1], then there is an *s*-*t* path *P* ^*\**^ which contains all the *α*-safe paths of *G*(*A, B*).

*Proof:* As defined above, for an edge *e* let *p*(*e*) ∈ [0, 1] denote the proportion of optimal paths it is part of. It is clear that

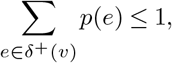

for all nodes *v*, where *δ*^+^(*v*) is the set of all outgoing edges of *v*.

We assume the opposite, meaning that there are two *α*-safe edges *e*_1_ and *e*_2_ for which no common *s*-*t* path exists. Since by definition of *α*-safe edges, *p*(*e*_1_) *>* 0.5 and *p*(*e*_2_) *>* 0.5, we obtain the contradiction

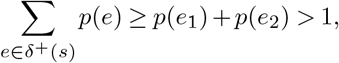

as the paths that cross *e*_1_ are distinct of the paths that cross *e*_2_ but all start at node *s*. Finally, since each pair of *α*-safe edges share a path, there must exist a single path *P* ^*\**^ containing all *α*-safe edges.

Next, we give an approach on how to find such a path *P* ^*\**^.

#### Lemma 2

Let *G* = (*V, E*) be a DAG with source *s* and sink *t*. Given a set of edges *A* = {*e*_1_, *e*_2_, …, *e*_*k*_ *}*, we can find an *s*-*t* path that either contains all the edges in *A* or return the information that such path does not exist, in time 𝒪 (|*V* | + |*E*|).

##### Algorithm 1

Pseudocode for inferring alignment-safety windows

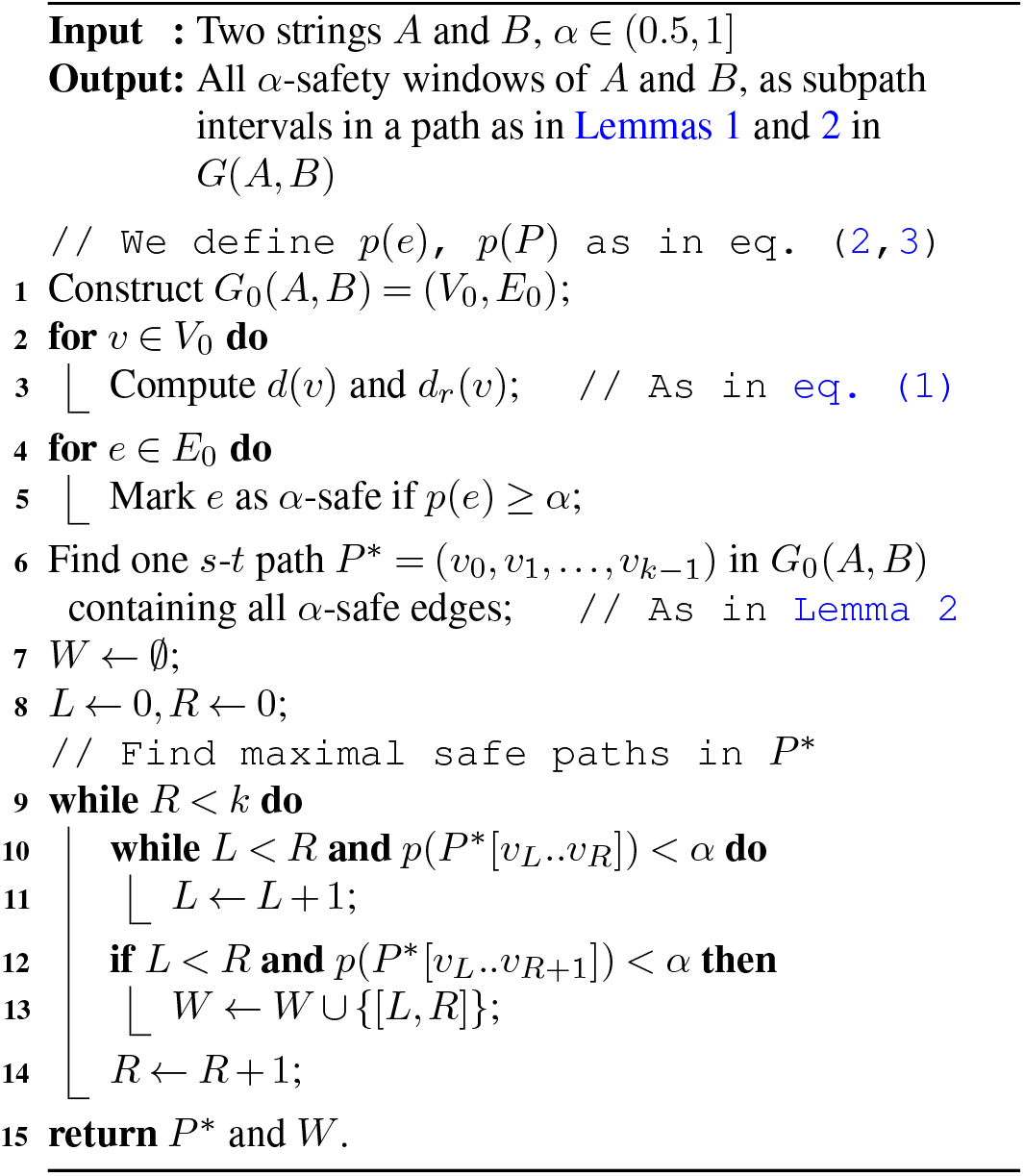

*Proof:* Let *T* be a topological order of all nodes in *V* and let *A* be sorted topologically by the tails of the edges. Assume we have a path from *s* to an edge *e*_*i*_ = (*u*_*i*_, *v*_*i*_). The task is to determine a path to the edge *e*_*i*+1_ = (*u*_*i*+1_, *v*_*i*+1_). For any path 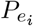 from *v*_*i*_ to *u*_*i*+1_ it must hold that *T*∈ [*x*] [*T* [*v*_*i*_], *T* [*u*_*i*+1_]] =: *I* (*⊆V*_0_) for all nodes *x* contained in 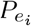. To achieve this, we perform a graph traversal (for example, depth-first search) in *I* and visit every node at most once. If *u*_*i*+1_ is not found after the graph search, it follows that the path does not exist. At the end of this procedure, if no edge in *A* is left, we just connect the path to *t*.

In Algorithm 1, we provide the pseudo-code to infer safety-windows. In lines 1 to 6, we construct the subgraph *G*_0_(*A, B*) of optimal paths, calculate the ratios *p*(*e*) for all *e* ∈ *E* and find a path *P* ^*\**^ according to Lemma 1. Next, we calculate the safety-windows with a two-pointer algorithm on *P* ^*\**^. The iterator variable here is *R* (right pointer), and we use a second left pointer *L* with *L* ≤*R*. The idea is to keep the value *L* as small as possible, to maintain the safety-windows left maximal, while moving *L* up until the interval [*L, R*] of *P* ^*\**^ becomes *α*-safe. On line 13 we add the interval into the list of safety-windows if it is also right-maximal (line 12).

#### Theorem 1

Let *α*∈ (0.5, 1]. Given two strings *A* and *B* of length *n* and *m*, respectively, Algorithm 1 computes the safety windows of *A* and *B* in time 𝒪 (*n· m*), assuming constant-time arithmetic operations.

*Proof:* We analyse the runtime line by line. First, constructing *G*_0_(*A, B*) can be performed in 𝒪 (*n ·m*) by the standard dynamic programming approach. As shown earlier, *d*(*v*) overall can be calculated in linear time (here 𝒪 (*n ·m*)) for DAGs. Line 5 runs in constant time, so this for-loop also runs in linear time.

Next, in line 6 we find an *s*-*t* path *P* ^*\**^ containing all *α*-safe edges, using the approach given in Lemma 2. Note that the path exists by Lemma 1.

Finally, we calculate the safety windows by iterating over *P* ^*\**^ and store them in the set *W*. Both variables *R* and *L* require at most 𝒪 (*n* + *m*) iterations and such *W* will at most contain 𝒪 (*n* + *m*) safety windows.

Let [*f, r*) ∈ *W*. According to the choice of *f* in line 10, such an interval is *α*-safe. It is maximal and thus a safety window, since neither [*f* −1, *r*) ∈ *W* nor [*f, r* + 1) ∈ *W*.

Next, we show that all safety windows are in *W*. Let [*f, r*) be a safety window. According to the iteration in line 9, *R* will eventually be equal to *r*. Consider the start of this iteration. If *L* ≤*f*, the iteration in line 10 will ensure that *L* will be equal to *f* by definition of safety windows and we insert [*f, r*) = [*L, R*) into *W*. We now assume *L > f*. In previous iterations *R*≤ *r*, we must have increased *L* to make it larger than *f*. But since [*f, r*) is *α*-safe, all intervals [*f, R*) for *R*≤ *r* are *α*-safe, and we never increase *L* if *L* = *f* and *R*≤ *r*.

Note that we assume that arithmetic operations run in constant time. If *G*_0_(*A, B*) contained all vertices in *G*(*A, B*), then the number of paths will be equal to the exponentially growing Delannoy numbers [Banderier and Schwer, 2005] (i.e. if *G*_0_(*A, B*) = *G*(*A, B*), then 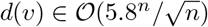, as this value is iteratively the sum of three previous values). Though, even if *G*_0_(*A, B*) only consists of nodes close to the diagonal line, the *d* values can grow up to 𝒪 (2*n*).

### Introducing gap penalties

So far, we only considered scores given by a substitution matrix *M* [1..*σ*][1..*σ*] for *σ* characters as well as by gaps through insertions and deletions when determining (sub)optimal global alignments between two protein sequences. While accounting for gaps in a global pairwise alignment is necessary to retrieve biologically relevant alignment configurations, finding the right balance between gap introduction and similarity optimisation is an important consideration to make.

To achieve this, we generalise the gap scoring function to be affine-linear. This means that when creating a gap of length *f* in the alignment, we obtain the score gap*p,g* (*f*) = *p* + *f ·g*. The value *p* ∈ *ℤ* is the *gap penalty*, chosen in order to minimise the number of gaps in an optimal alignment. Before having introduced a gap score, we have set *p* = 0. To accommodate gap scores in our previously defined DAG *G*, we can slightly modify the graph to address the known issue that paths are then not in 1:1 correspondence to alignments anymore. Following Myers and Miller [1988], we replace each node (*i, j*) by three nodes *C*(*i, j*), *D*(*i, j*) and *I*(*i, j*). Nodes *D* and *I* correspond to being inside a gap, erasing the characters in *A* and *B* respectively, while node *C* is the default node, from which we can decide to either match two characters or to introduce a new gap. We thus create the following edges for *C*: *C*(*i, j*) → *C*(*i* + 1, *j* + 1) to match *A*_*i*_ with *B*_*j*_, *C*(*i, j*) → *D*(*i* + 1, *j*) to align *A*_*i*_ with a gap and *C*(*i, j*) →*I*(*i, j* + 1) to align *B*_*j*_ with a gap. The last two of the edges also define the start of a gap. For *D* (and analogously for *I*), we create the following edges: *D*(*i, j*) →*C*(*i, j*) to close a gap and *D*(*i, j*) →*D*(*i* + 1, *j*) to extend a gap. The new graph is an *s*-*t* DAG of size 𝒪 (*n ·m*) which retains all properties to calculate safety-windows as previously described. Additionally to the substitution matrix and the gap score, we assign edge weights for closing and opening a gap, commonly chosen as 0 for closing and some negative integer for opening in order to punish creating too many gaps.

### Extending our DAG approach to incorporate suboptimal alignments

We previously stated that the graph *G*_0_(*A, B*) has the property that all its *s*-*t* paths are optimal paths in *G*(*A, B*). This gives us a compact data structure able to store exponentially many optimal paths. We now generalise the subgraph *G*_0_ to contain also suboptimal paths. This is achieved by introducing a critical threshold Δ, which users can specify to either extend or narrow down the suboptimal alignment space. For a given Δ≥ 0, an *s*-*t* path in *G*(*A, B*) is said to be Δ*-suboptimal* if it is of weight at least *OPT* −Δ, where *OPT* denotes the weight of an optimal path. We denote paths that appear in at least an *α*-proportion of all Δ- suboptimal paths in *G*(*A, B*) as (*α*, Δ)*-safe* and define maximal (*α*, Δ)-safe paths as (*α*, Δ)*-safety windows*.

Following the notion of Naor and Brutlag [1994], we define *G*_Δ_(*A, B*) = (*V*_Δ_, *E*_Δ_) to be the subgraph of *G*(*A, B*) which is induced by the set of edges *e*∈ *E* such that there is a Δ- suboptimal *s*-*t* path in *G*(*A, B*) crossing *e*. By definition, *G*_Δ_(*A, B*) contains all Δ-suboptimal *s*-*t* paths in *G*(*A, B*). Additionally, however, it may contain *s*-*t* paths that are not Δ-suboptimal. This implies that some paths of *G*_Δ_(*A, B*) appear in at least a proportion *α* of *s*-*t* paths which are not necessarily (*α*, Δ)-safe paths (in *G*(*A, B*)).

Naor and Brutlag [1994] have shown that *G*_Δ_(*A, B*) is the *smallest* subgraph of *G*(*A, B*) that contains all Δ-suboptimal *s*-*t* paths. In fact, our data spanning tree-of-life scale protein sequence diversity does not indicate that these spurious non-suboptimal paths have any effect in practice. Naor and Brutlag [1994] argue that the only possible solution to capture only suboptimal paths would be to naively enumerate over all suboptimal paths, which is unfeasible, since in the worst case there can be exponentially many suboptimal paths of size *G*(*A, B*). We address this limitation by approximating (*α*, Δ)-safe paths with those paths of *G*_Δ_(*A, B*) appearing in at least a proportion *α* within its set of *s*-*t* paths. We compute these paths according to Algorithm 1, with the only difference that we start with *G*_Δ_(*A, B*) in line 1 instead of *G*_0_(*A, B*).

To compute *G*_Δ_(*A, B*), we proceed as in [Naor and Brutlag, 1994]: For all nodes *v* ∈ *V*, we find the weight *w*(*v*) of an optimal *v*-*t* path of size *G*(*A, B*) in linear time by traversing the nodes in reverse topological order, such that the weights of all out-neighbours of *v* are computed when we reach *v*. Similarly, we compute the weight *w*^*r*^(*v*) of optimal *s*-*v* paths for all *v* ∈ *V*. An edge *e* = (*u, v*) ∈ *E* is now part of *G*_Δ_(*A, B*) only if *w*^*r*^(*u*) + *w*(*v*) + *w*(*e*) ≥ *OPT* −Δ, where *w*(*e*) denotes the weight of the edge *e*. We can thus construct *G*_Δ_(*A, B*) in linear time and proportional to the size of *G*(*A, B*).

### Implementing Alignment-Safety into the scientific software EMERALD

We designed EMERALD to efficiently implement our theoretical methodology to explore the suboptimal alignment space by inferring alignment-safe intervals when performing pairwise protein sequence comparisons. In detail, EMERALD accepts a protein sequence cluster in FASTA format of *k* sequences as input and returns *k*− 1 safety-window sets in a custom safety window format. Our tool aligns all sequences in the cluster against a user-selected representative sequence of the cluster, which by default is the first sequence in the FASTA file, thereby maximising the score of the alignment. EMERALD by default uses BLOSUM62 as substitution matrix, however, any affine-linear gap cost function can be used via the command line parameters. This makes EMERALD extendable to sequence alignments beyond protein sequences. For a cluster containing *k* sequences, the goal is to be able to compare the safety-windows of all *k*− 1 pairs with each other. Thanks to Lemma 1, we can project the safety windows to the sequences. Node indices in the alignment graph are of the form (*i, j*) with 0 ≤*i* ≤*n* and 0 ≤*j* ≤*m*, and such safety windows can be written as [(*i*_1_, *j*_1_), (*i*_2_, *j*_2_)]. For example, given the strings *S* = “*AB*” and *T* = “*BC*” and the alignment “*AB* − “ and “ −*BC*”, a safety window of the first gap would be of the form [(0, 0), (1, 0)]. Even though by usual convention *T* [0] = *B*, this position is not part of the safety window. EMERALD then returns the interval [0, 1] for string *S* and [0, 0] for string *T*. In other words, the tuple indices of the nodes are *between* the string characters. This coincides with the fact that the node set is the product {0, …, *n}× {*0, …, *m}* (i.e. (*n* +1) *·* (*m* +1) nodes) and with the fact that we want to be able to include gaps in safety windows.

## Data Access

EMERALD’s source code [Grigorjew et al., 2023] can be accessed at https://github.com/algbio/emerald. All data preprocessing is implemented as a reproducible and parametrised Snakemake pipeline [Mölder et al., 2021] and computationally reproducible scripts can be found at https://github.com/algbio/emerald-analysis. All datasets, including the 396k protein sequences, DIAMOND-clusters, EMERALD-output, and annotation files can be retrieved from figshare: https://doi.org/10.6084/m9.figshare.21720299.

## ACKNOWLEDGEMENTS

This work was partially funded by the European Research Council (ERC) under the European Union’s Horizon 2020 research and innovation programme (grant agreement No. 851093, SAFEBIO), and partially by the Academy of Finland (grants No. 322595, 328877, 352821). We thank V. Alva, H. Ashkenazy, C. Behr, A. Dobbelstein, F. Gabler, Lukas Maischak, K. K. Ullrich and D. Weigel for their interest and helpful discussions. H.G.D. and B.B. thank Detlef Weigel for ongoing support and sponsorship of their work. H.G.D. thanks the BMBF-funded de.NBI Cloud within the German Network for Bioinformatics Infrastructure (de.NBI) (031A532B, 031A533A, 031A533B, 031A534A, 031A535A, 031A537A, 031A537B, 031A537C, 031A537D, 031A538A) for support. Finally, H.G.D. and B.B. acknowledge that their work was supported by the Max Planck Society.

## Author contributions

A.T. and H.G.D. designed and led this study. An.G. implemented EMERALD with feedback from Ar.G., F.D., B.B., A.T., and H.G.D. Data analysis was performed by An.G., Ar.G., F.D., B.B. Data interpretation was performed by A.T., H.G.D., An.G., Ar.G., F.D., and B.B. The manuscript was written by H.G.D., An.G., and A.T., with contributions from Ar.G., F.D., B.B.

---

**Supplementary Table 1:** Summary statistics of pairwise sequence assessments in regard to their parameter-specific safety-window length construction. Shown are the arithmetic means of aligned sequence lengths and their respective safety window counts and average safety-window lengths for ∼400k SwissProt sequences. In this assessment, we also included the previously removed ∼4k sequences which did not include any stable bases. We observe that when increasing the Δ parameter, this leads to increases in the number of safety windows while decreasing their average length. On the other hand, when increasing *α*, we observe a decrease in the number of safety windows and an average increase of safety-window lengths.

| Parameters | ID range | Number of | | Average length of | | Proportion of safety windows of length $\geq 50\%$ |
| --- | --- | --- | --- | --- | --- | --- |
|  |  | sequences | safety windows | sequences | safety windows |  |
| $\alpha = 0.51, \Delta = 8$ | [0%, 100%] | 400408 | 2882661 | 321.89 | 40.49 | 6.9% |
| $\alpha = 0.75, \Delta = 0$ | [0%, 100%] | 400408 | 1059216 | 321.89 | 117.96 | 30.5% |
|  | [0%, 20%] | 2922 | 16674 | 467.74 | 75.58 | 5.3% |
|  | [20%, 40%] | 157236 | 551251 | 339.47 | 92.50 | 18.9% |
|  | [40%, 70%] | 197051 | 433730 | 308.90 | 137.17 | 40.4% |
|  | [70%, 100%] | 44674 | 61172 | 304.80 | 221.11 | 71.7% |
| $\alpha = 0.75, \Delta = 2$ | [0%, 100%] | 400408 | 2075297 | 321.89 | 56.67 | 10.5% |
| $\alpha = 0.75, \Delta = 4$ | [0%, 100%] | 400408 | 2375169 | 321.89 | 47.87 | 8.3% |
| $\alpha = 0.75, \Delta = 6$ | [0%, 100%] | 400408 | 2532554 | 321.89 | 43.66 | 7.4% |
| $\alpha = 0.75, \Delta = 8$ | [0%, 100%] | 400408 | 2643650 | 321.89 | 40.73 | 6.8% |
|  | [0%, 20%] | 2922 | 52110 | 467.74 | 14.70 | 0.03% |
|  | [20%, 40%] | 157236 | 1522093 | 339.47 | 26.27 | 1.7% |
|  | [40%, 70%] | 197051 | 989107 | 308.90 | 54.51 | 11.3% |
|  | [70%, 100%] | 44674 | 88974 | 304.80 | 149.21 | 47.5% |
| $\alpha = 0.75, \Delta = 10$ | [0%, 100%] | 400408 | 2784429 | 321.89 | 37.67 | 6.1% |
| $\alpha = 1.00, \Delta = 8$ | [0%, 100%] | 400408 | 1844829 | 321.89 | 53.41 | 9.4% |

**Supplementary Figure 1:**
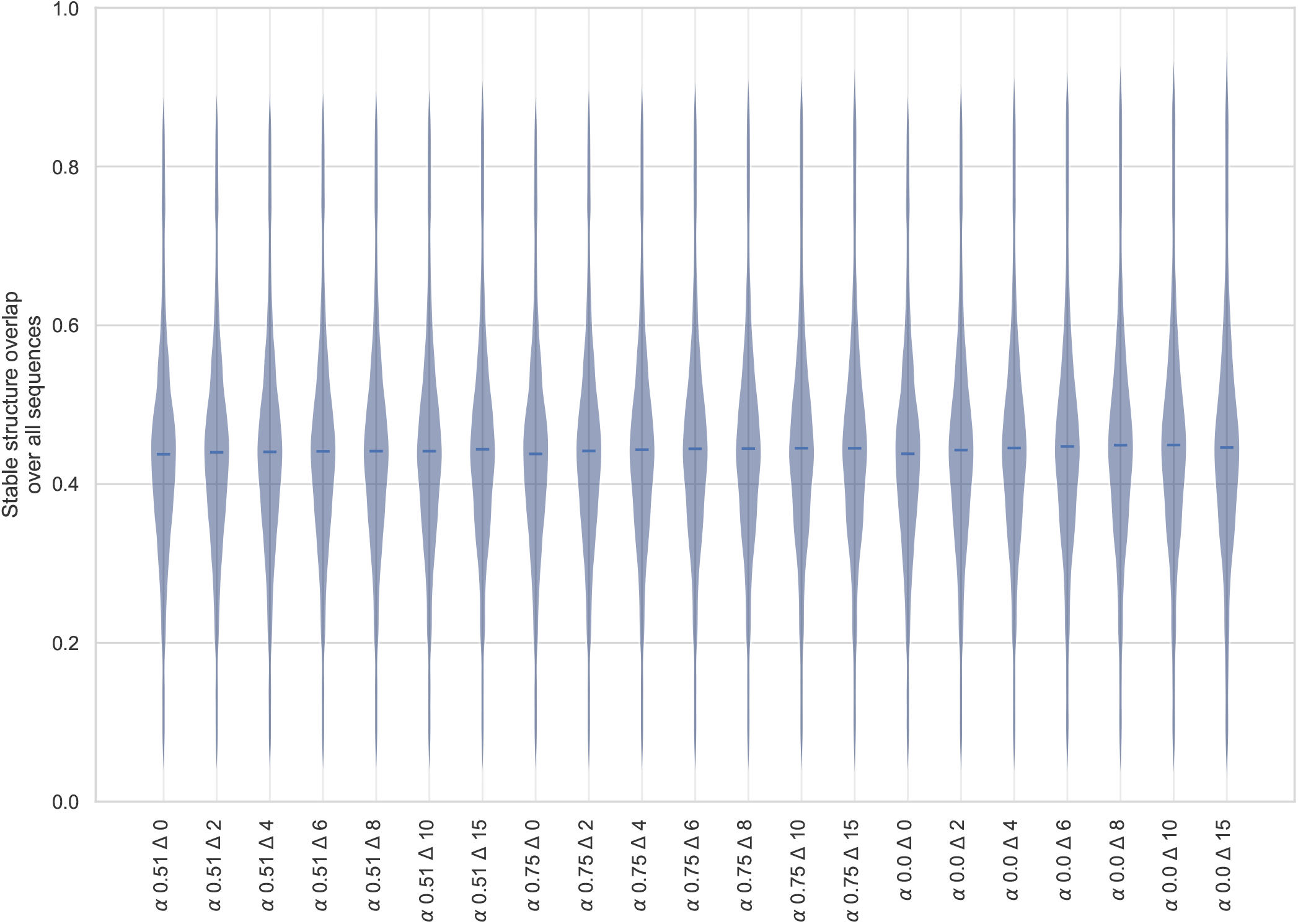
Stable structure overlap for each pair of *α* = 0.51, 0.75, 1 and Δ = 0, 2, 4, 6, 8, 10, 15 on the whole dataset of 396k sequences. 16 pairs of aligned sequences had to be filtered out due to a safety coverage of 0. The stable structure overlap is throughout all pairs constant with the median at 43%. This is exactly the proportion of stable positions within *α*-safety.

**Supplementary Figure 2:**
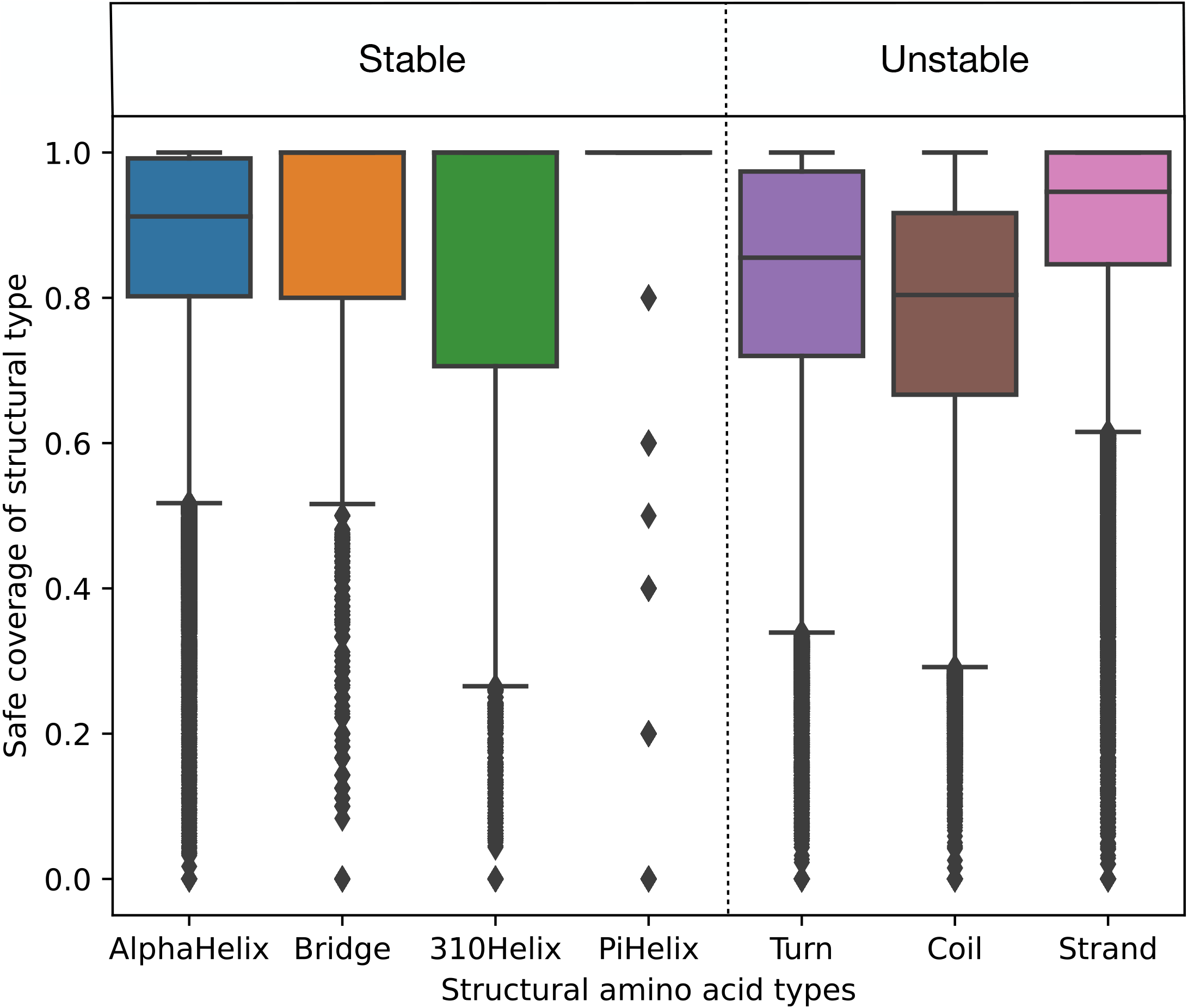
Safety coverage restricted on each structural type as assigned by STRIDE (i.e. for every structural type *A* (number of amino acids of type *A* that are safe) / (total number of amino acids of type *A*)) with EMERALD parameters *α* = 0.75 and Δ = 8, throughout all 400k sequences, including previously removed 4k sequences which did not include any stable amino acids. Every point is the safety coverage restricted to the corresponding structure type for all safety windows of a sequence. In the experimental results, we defined Turn, Coil and Strand as unstable, and all other structural types as stable. For example, AlphaHelix has a median safety coverage of 90% throughout all sequences. We observe that stable amino acids have a larger safety coverage than unstable ones, with the exception of Strand having a higher coverage than AlphaHelix. The distribution of the structural types throughout all sequences is as follows, rounded to one decimal place: AlphaHelix: 40.3%, Bridge: 0.8%, 310Helix: 3.2%, PiHelix: 0.007%, Turn: 17.7%, Coil: 19.1% and Strand: 18.7%.

**Supplementary Figure 3:**
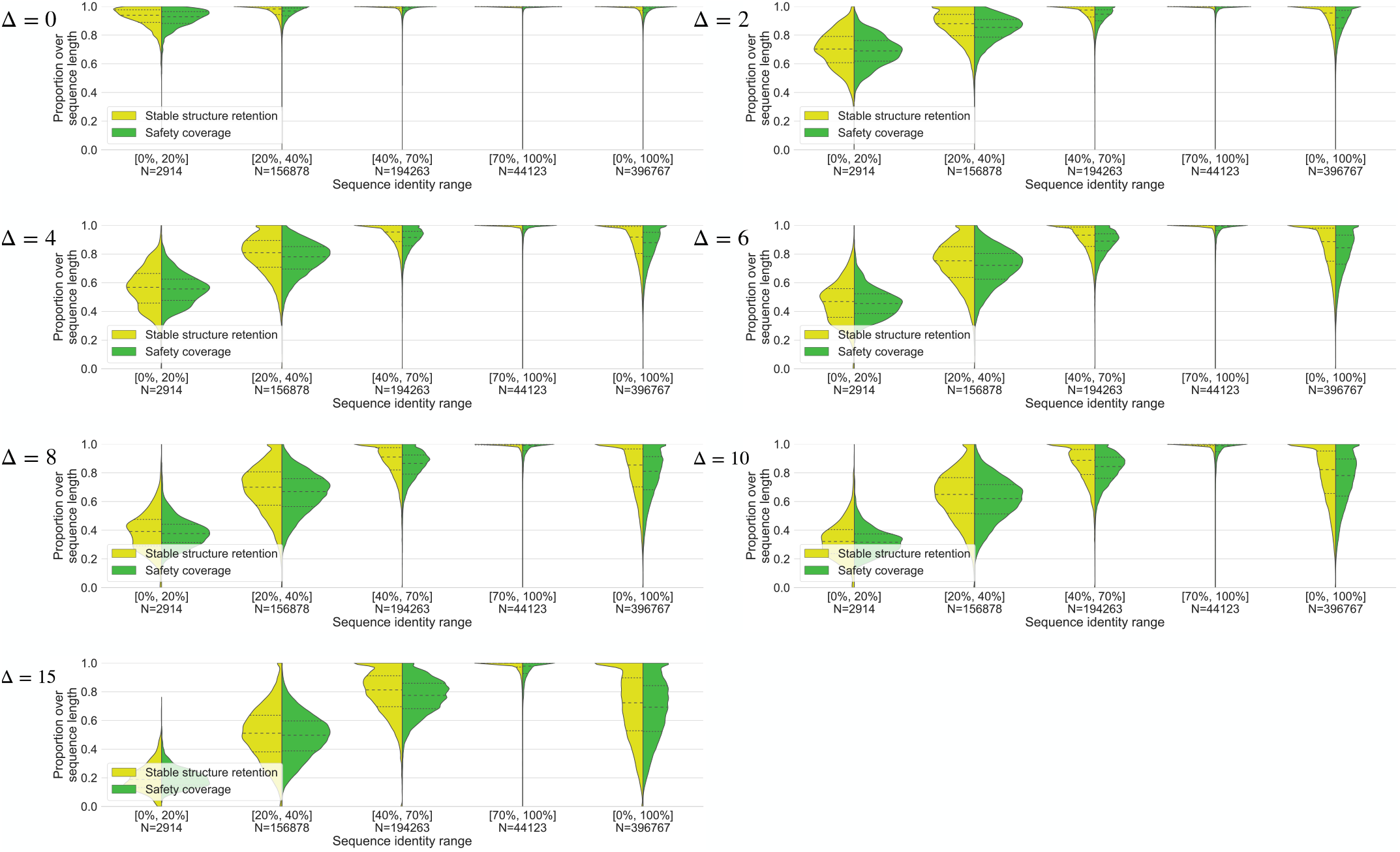
Stable structure retention compared to safety coverage for *α* = 1, Δ = 0, 2, 4, 6, 8, 10, 15 and several identity ranges. The median and the quartiles are indicated in black.

**Supplementary Figure 4:**
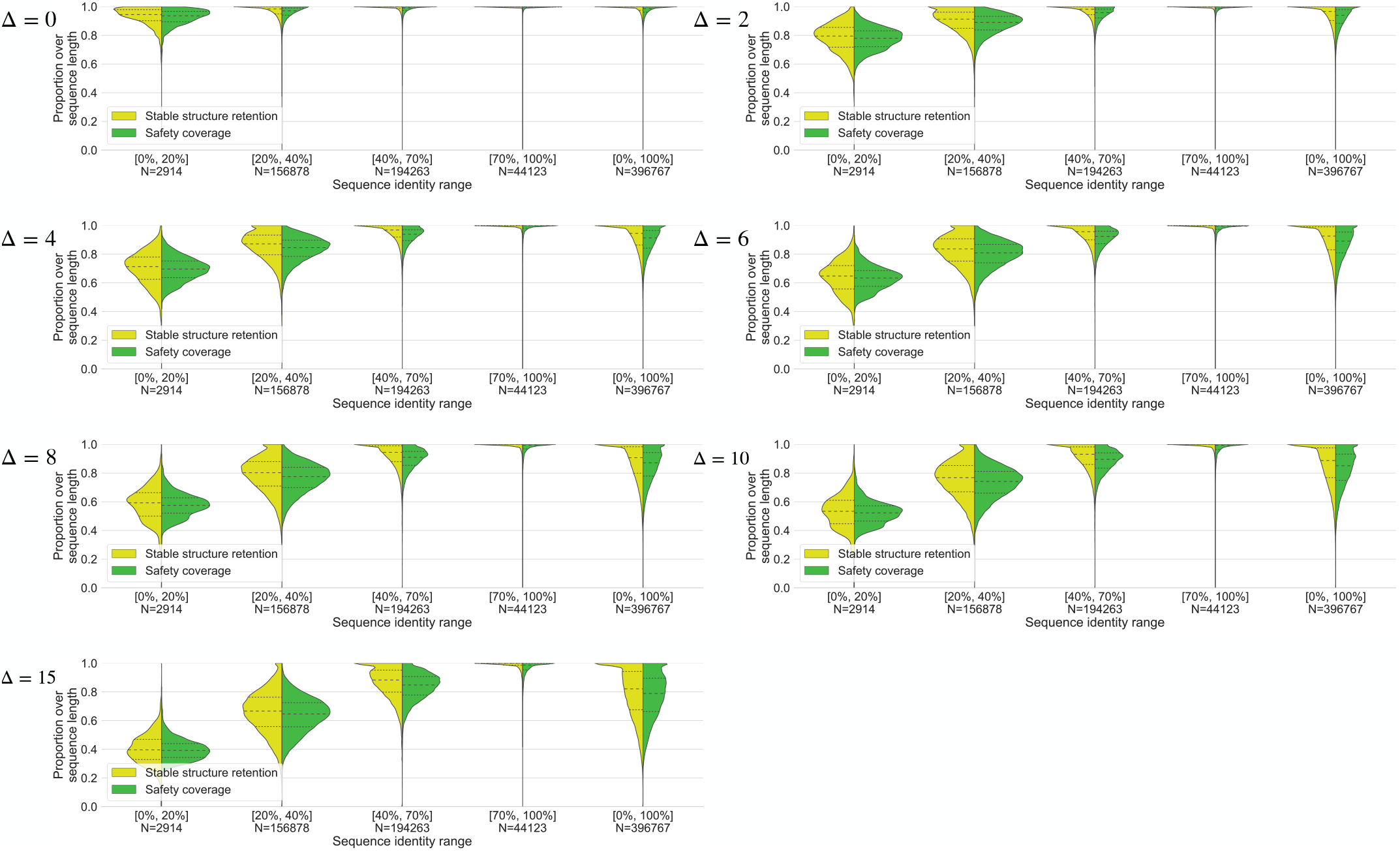
Stable structure retention compared to safety coverage for *α* = 0.75, Δ = 0, 2, 4, 6, 8, 10, 15 and several identity ranges. The median and the quartiles are indicated in black.

**Supplementary Figure 5:**
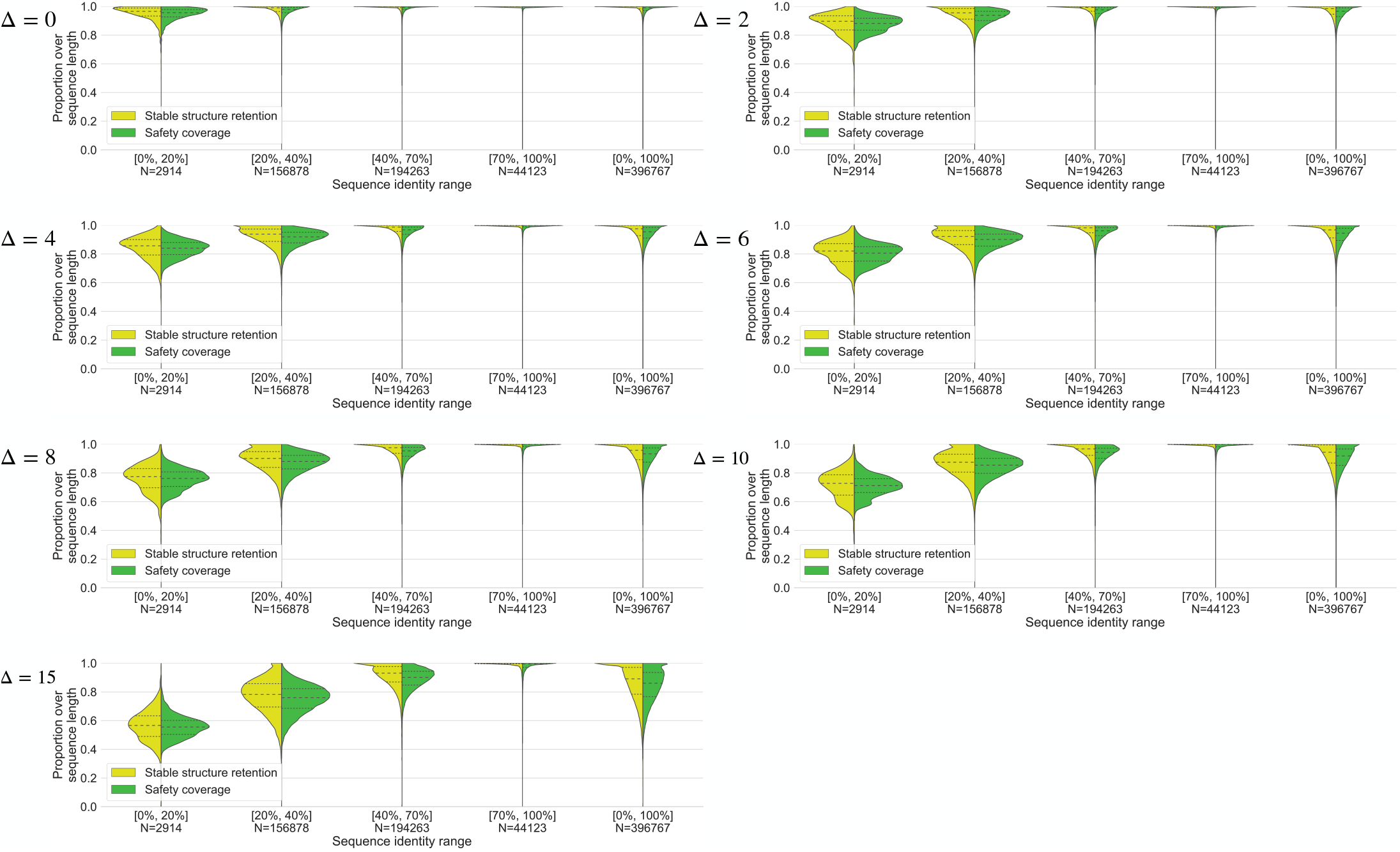
Stable structure retention compared to safety coverage for *α* = 0.51, Δ = 0, 2, 4, 6, 8, 10, 15 and several identity ranges. The median and the quartiles are indicated in black.

**Supplementary Figure 6:**
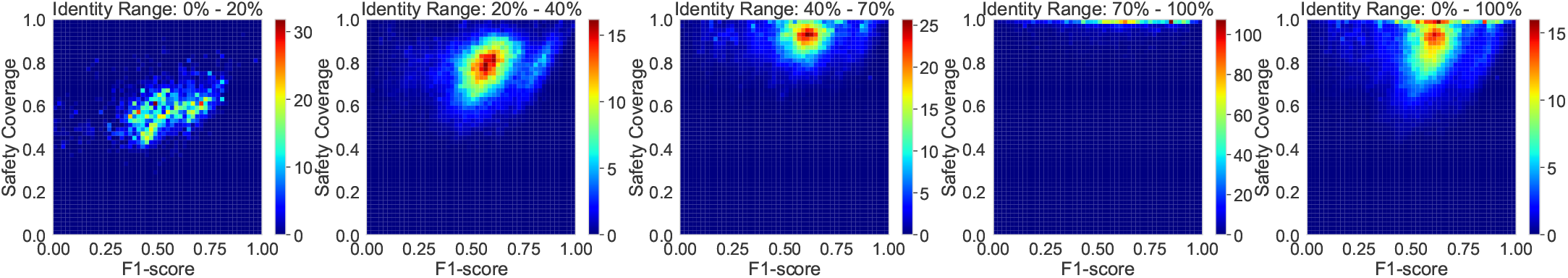
F1-score for *α* = 0.75, Δ = 8, for sequences in clusters of varying identity ranges. We can see that the F1-score lies consistently in a range around 50%, while in the lower cluster identities the sequence is covered less by safety windows. This indicates that EMERALD performs better in the identity range 20-70%.

**Supplementary Figure 7:**
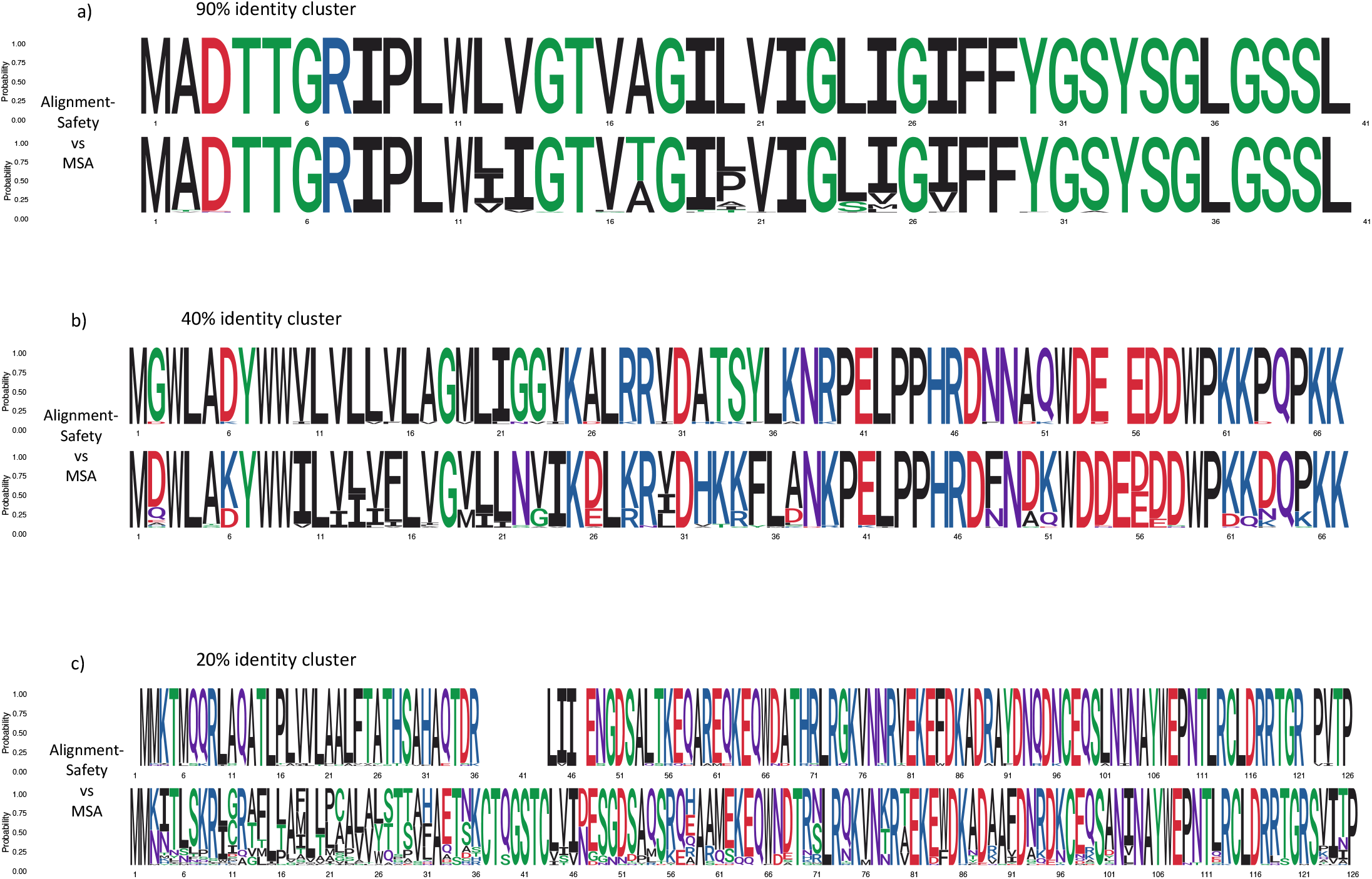
Direct comparison of sequence logos between multiple sequence alignments (MSAs) and one-vs-all (cluster members against cluster representative) EMERALD alignments derived from one DIAMOND DeepClust clusters with various identity ranges (90% identity, 40% identity, 20% identity). For MSAs, the sequence logos are generated directly from the MSA output, while for EMERALD alignments sequence logos are generated via interpolation of the conserved safety windows among different pairwise alignments (cluster representative vs cluster members). In the x-axis, the position of each amino acid is represented, while the y-axis has the probabilities defined by the relative frequencies of residue occurrence. Results are shown for a) (90% cluster identity), b) (40% cluster identity) and c) (20% cluster identity), the top sequence logos denote EMERALD derived alignments and the bottom sequence logo are generated from MSAs calculated using MUSCLE Edgar [2004]. Lower identity clusters contain larger proportions of biodiverse protein sequences. With increasing sequence diversity and lower identity boundaries in a cluster, the multiple sequence aligner attempts to optimally place alignment configurations such that the alignment of common regions is favoured. In contrast, sequence logos created with EMERALD show only the common regions that are intrinsically present throughout all pairwise alignments (or most alignments for *α* < 1) and by the definition of safety, it is guaranteed that such a conserved region is present in at least *α* suboptimal alignments. Overall, this figure illustrates that conserved regions obtained from MSAs and EMERALD increasingly differ with decreasing identity boundaries, thereby presenting opportunities for subsequent studies to explore the differences in biological interpretations when comparing information derived from the suboptimal alignment space (alignment-safety) and information derived from MSAs.

